# Single-cell transcriptomic atlas reveals increased regeneration in diseased human inner ears

**DOI:** 10.1101/2022.10.29.514378

**Authors:** Tian Wang, Angela H. Ling, Sara E. Billings, Davood K. Hosseini, Yona Vaisbuch, Grace S. Kim, Patrick J. Atkinson, Zahra N. Sayyid, Ksenia A. Aaron, Dhananjay Wagh, Nicole Pham, Mirko Scheibinger, Akira Ishiyama, Peter Santa Maria, Nikolas H. Blevins, Robert K. Jackler, Stefan Heller, Ivan A. Lopez, Nicolas Grillet, Taha A. Jan, Alan G. Cheng

## Abstract

Mammalian inner ear hair cell loss leads to permanent hearing and balance dysfunction. In contrast to the cochlea, vestibular hair cells of the murine utricle have some regenerative capacity. Whether human utricular hair cells regenerate remains unknown. Here we procured live, mature utricles from organ donors and vestibular schwannoma patients, and present a validated single-cell transcriptomic atlas at unprecedented resolution. We describe previously unknown markers of 25 sensory and non-sensory cell types, with genes of hair cell and supporting cell subtypes displaying striking divergence between mice and humans. We further uncovered transcriptomes unique to hair cell precursors, which we validated to be 14-fold more robust in vestibular schwannoma utricles, representing ongoing regeneration in humans. Lastly, trajectory analysis of the supporting cell-hair cell axis revealed 5 distinct patterns of dynamic gene expression and associated pathways, including mTOR signaling and synaptogenesis. Our dataset constitutes a foundational resource, accessible via a web-based interface, serving to advance knowledge of the normal and diseased human inner ears and tools to stimulate human inner ear regeneration.

## Introduction

Hearing and vestibular disorders affect over 460 million people worldwide (WHO, 2021). In the inner ear, auditory and vestibular hair cells are essential for detecting sound and head motions. Genetic mutations, ototoxins, noise and aging are known to cause hair cell degeneration, leading to permanent hearing loss and vestibular dysfunction (Iyer and Groves, 2021; Maudoux et al., 2022; Morton and Nance, 2006). While the loss of mammalian auditory hair cells is irreversible, mouse models have shown that vestibular hair cells turn over during homeostasis and regenerate after damage (Bucks et al., 2017; Burns et al., 2012; Forge et al., 1993; Golub et al., 2012; Kawamoto et al., 2009; Sayyid et al., 2019; Wang et al., 2015; Wang et al., 2019). Whether these phenomena similarly occur in the human vestibular system has not yet been demonstrated, in part because obtaining normal and diseased human inner ear tissues is exceedingly challenging.

The human inner ear is housed in the temporal bone, making it accessible only via surgery (Feghali and Kantrowitz, 1994; Oghalai et al., 2000). Because unroofing the human inner ear renders it non-functional, the majority of work done on the human inner ear has been performed on postmortem, cadaveric tissues (Hizli et al., 2016; Kaya et al., 2017; Lopez et al., 2005; Lopez et al., 2016a; Merchant, 1999; Merchant et al., 2000; Oghalai et al., 2000; Sans et al., 1996; Senn et al., 2020; Watanuki and Schuknecht, 1976). A few studies on viable inner ear tissues from fetuses and organoids derived from embryonic or induced pluripotent stem cells have characterized morphological and molecular features of the developing human inner ear (Chen et al., 2007; Chen et al., 2009; Johnson Chacko et al., 2019; Nie and Hashino, 2020; Roccio et al., 2018; Ueda et al., 2021), however, the transcriptomic signature of the healthy, adult human inner ear has remained elusive because of the difficulty of obtaining live samples for such experiments.

An alternative approach is to analyze live inner ear tissues extracted during surgery from patients affected by vestibular schwannoma, a benign tumor of the myelin-forming cells of the vestibulocochlear nerve. Patients with vestibular schwannoma develop hearing loss and vestibular dysfunction (Gupta et al., 2020; Matthies and Samii, 1997), presumably due to spiral ganglion cell loss caused by nerve compression (Eggink et al., 2022) or hair cell degeneration caused by secreted exosomes (Soares et al., 2016). In certain cases, vestibular schwannoma patients warrant surgical resection via a translabyrinthine approach where inner ear tissues are sacrificed (Feghali and Kantrowitz, 1994; House, 1964). Utricles harvested from vestibular schwannoma patients during surgery have served as the primary model system to characterize hair cell morphology (Hizli et al., 2016; Sans et al., 1996; Taylor et al., 2015), their capacity to regenerate after aminoglycoside damage *in vitro* (Taylor et al., 2018; Warchol et al., 1993), and responsiveness to Atoh1-overexpression to induce ectopic hair cells (Taylor et al. 2018). However, it remains undetermined in these diseased samples whether hair cell regeneration occurs spontaneously in human utricles *in vivo*.

In this study, we successfully sequenced and validated utricular single-cell transcriptomes in adult organ donors and vestibular schwannoma patients Our computational analysis showed that adult human utricles of healthy and schwannoma patients are composed of 25 cell types with unique molecular signatures. Gene expression of human hair cells and supporting cells were well conserved relative to those in mice, with divergent features among hair cell and supporting cell subtypes. Single-cell transcriptomic trajectory analysis revealed dynamic changes as supporting cells transition into hair cells in both vestibular schwannoma and organ donor utricles. As predicted in tissues, hair cell loss was increased in schwannoma utricular samples compared to ones from organ donors and cadavers. Finally, we observed a ~14-fold increase of hair cell precursors (POU4F3^+^ and GFI1^+^) in vestibular schwannoma compared to organ donor utricles. Together, we have demonstrated spontaneous, unmanipulated hair cell regeneration in the adult human utricle and the underlying transcriptional changes during homeostasis and in response to damage. These rare datasets represent foundational tools to guide the investigation of human hair cell regeneration and define molecular differences between vestibular organs in humans and other mammalian species.

## Results

### Transcriptional heterogeneity of the mature human utricle

To characterize the human utricle, we harvested tissues from two cohorts: organ donors and vestibular schwannoma patients (Table S1). We previously devised a surgical approach to procure unilateral or bilateral utricles from organ donors (OD) (Aaron et al., 2022; Vaisbuch et al., 2022), who lacked any history of auditory or vestibular dysfunction (Table S1). In parallel, utricles were procured from patients with unilateral vestibular schwannomas and ipsilateral auditory and/or vestibular deficits, undergoing a translabyrinthine approach for tumor resection.

Because vestibular schwannoma patients suffer from auditory and/or vestibular deficits and because hair cell degeneration is detected in their cochlear and utricular organs postmortem (Hizli et al., 2016; Sans et al., 1996), we hypothesized that vestibular schwannoma utricles are damaged whereas those from organ donors may serve as normal controls (Figure 1A, Figure S1A, Table S1). Two vestibular schwannoma and one organ donor utricles were subject to single-cell isolation and sequencing. Sensory epithelia were enzymatically and mechanically purified from surrounding stromal and neural tissues (Figure S1B), yielding a total of 12,765 single cells that were sequenced (7,376 organ donor cells and 5,389 vestibular schwannoma cells, Figure 1B’, Figure S2A). Samples from four captures (two vestibular schwannoma samples with one capture each, and one organ donor sample in two captures) were sequenced (10x Genomics platform), and cells underwent quality control metrics and downstream analyses (Figure S1C). Our integrated dataset (both organ donor and vestibular schwannoma) contained a median of 2,061 genes per cell (Figure S1D) and unsupervised clustering identified 25 cell states (Figure 1B). Similarity weighted non-negative embedding (SWNE) was used for all 2-dimensional visualization of reduced dimensions to optimally capture global and local relationships among cells and clusters (Parekh et al., 2018). We annotated the cell clusters based on known marker genes in the mammalian inner ear (Figure 1B, Figure S1E), including 25 different cell types with 16,866 cluster-defining genes (Wilcoxon test, FDR <0.01, Figure 1C, Table S2): supporting cells (*GFAP*, *SNORC*, *SFRP2*, clusters 0, 2, 4, 5, 7, 9 and 10), hair cells (*CIB3*, *OTOF*, *MYO7A*, *TMC1*, clusters 13 and 17), hair cell precursors (*COX7A1, S100A6, DNAJB1,* cluster 20), roof cells (*CITED1*, cluster 18), dark cells (*BSND*, *KCNQ1*, *IGF1*, clusters 1, 6, 14 and 16), stromal cells (*PDGFRA*, clusters 3, 11 and 12), melanocytes (*PMEL*, *DCT*, *MLANA*, cluster 15), vascular cells (*FLT1*, cluster 24), macrophages (*CCL3*, clusters 19 and 21), pericytes (*RGS5*, *EDNRA*, cluster 22), and immune cells (*SERPINE2*, cluster 8 and 23).

**Figure 1.**
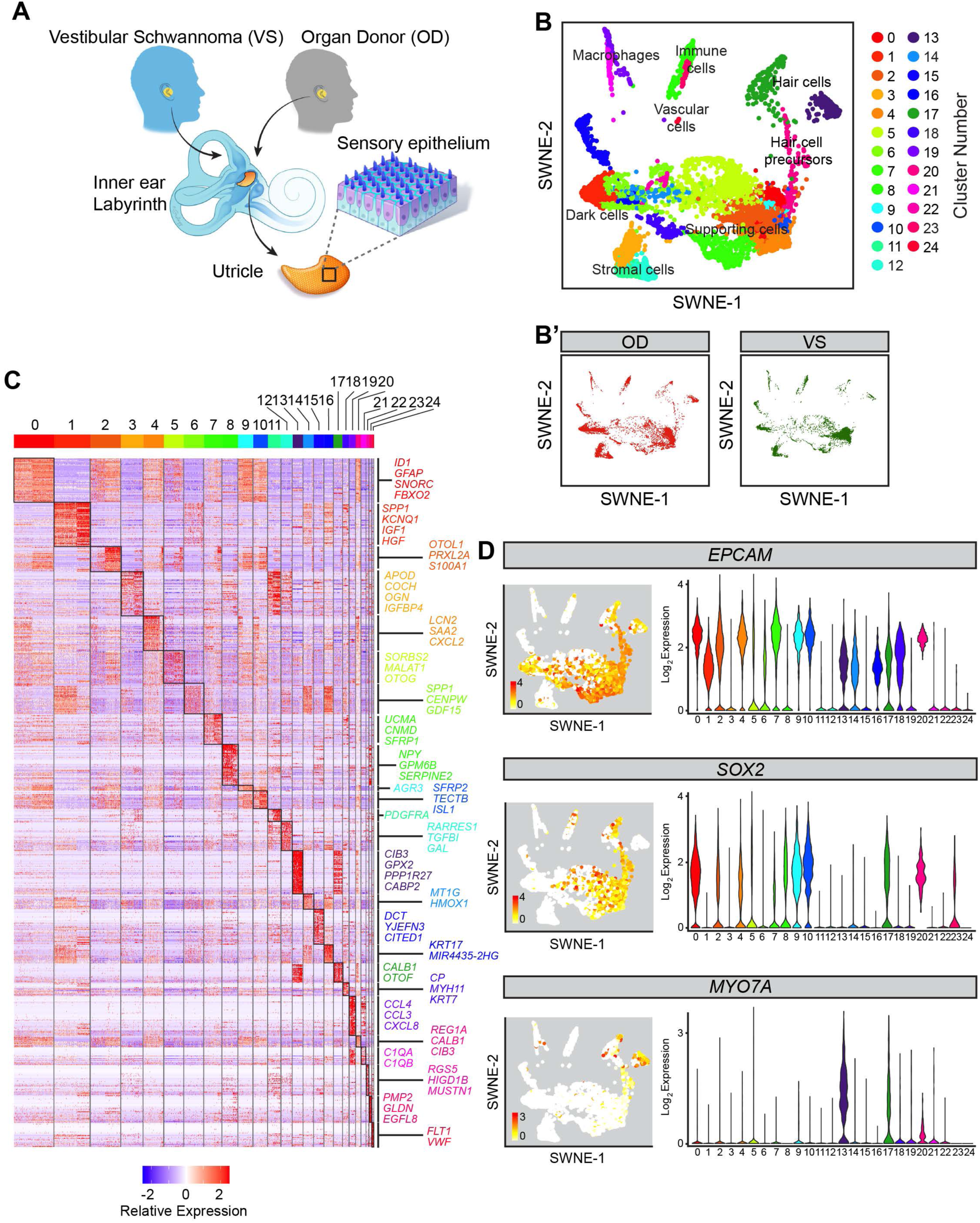
Sensory and non-sensory cell types in utricles from vestibular schwannoma and organ donor patients. (A) Cartoon depicting procurement of human utricles from patients undergoing translabyrinthine resection of vestibular schwannoma (VS) and organ donors (OD). Samples from both sources were used for histological and single-cell RNA sequencing analyses. (B) SWNE plot of integrated dataset of two VS utricles and one OD utricle showing 12,765 single cells following all quality control steps including exclusion of doublets. Twenty-five cell clusters were identified, and marker genes were used to annotate the sensory and non-sensory cell types. (B’) The SWNE plot from (B) is decomposed into the cells originating from OD and VS subjects. There were 7,376 and 5,389 cells from the OD and VS samples, respectively. (C) Heatmap showing differentially expressed genes among the 25 cell clusters. The top differentially expressed genes of each cluster are shown on the right side of the heatmap. A full list of these genes is found in Table S2. The heatmap is colored by relative expression from −2 (blue), 0 (white), and 2 (red). (D) Expression of established markers of epithelial cells (*EPCAM*), non-sensory cells and type II hair cells (*SOX2*) and sensory cells (*MYO7A*) within the cell clusters. SWNE plots colored by log_2_ expression (values of 0 in white to maximum in red as indicated) and violin plots colored by cell cluster are shown. *EPCAM* was highly expressed in clusters 0, 1, 2, 4, 5-7, 9, 10, 13, 14, 16-18, and 20; *SOX2* in clusters 0, 2, 4, 5, 7-10, 17, 20 and 23; and *MYO7A* in clusters 13 and 17, which represents hair cells. *SOX2* labels a subgroup of the *MYO7A^+^* cells indicating type II hair cells.

To further restrict our analyses to cells in the sensory epithelium, we focused on clusters defined as hair cells, hair cell precursors, and supporting cells (Figure S2A). As expected, *EPCAM* expression was robust in these sensory epithelial cell clusters (Figure 1D). Expression of *SOX2*, a known marker of supporting cells and type II hair cells in human, was enriched in a subset of hair cells and supporting cells, while expression of *MYO7A*, a known human generic hair cell marker (Taylor et al., 2015), is the highest in clusters 13 and 17 (Figure 1D). Sub-clustering and visualization with SWNE showed ten cell states representing utricle sensory epithelial cells (Figure S2A’). The final analysis consisted of 3,292 organ donor and 3,341 vestibular schwannoma utricular cells (Figure S2A’ and C). Differential gene expression analysis identified 5,694 cluster defining genes (Figure S2D, Table S3), yielding seven subsets of supporting cells (n=5,786 cells, clusters 0, 2, 4, 5, 7, 9 and 10), one putative hair cell precursor group (n=165 cells, cluster 20), and two hair cell subtypes (n=373 Type I hair cells, cluster 13 and n=309 Type II hair cells, cluster 17) (Figure S2B). Together, this dataset represents the molecular identities of distinct cell types of the adult human utricle.

### Molecular identity of human hair cells and supporting cells

The sensory epithelium is comprised of sensory hair cells and non-sensory supporting cells (Figure 2A). To identify hair cell- and supporting cell-defining genes, we performed a direct comparison without the putative hair cell precursors (Figure 2B). We identified 1,163 enriched genes in hair cells and 404 enriched genes in supporting cells with at least 2-fold expression difference (LFC>2, FDR<0.01, Figure 2C, Table S4). Among the 24 most differentially expressed genes in hair cells (*CABP2*, *CIB3, <u>GPX2</u>, ESPN, <u>CD164L2</u>, SYT14, <u>SNCG</u>, <u>CPE</u>, STRC, <u>SMPX</u>, <u>ABCA5</u>, POU4F3*) and supporting cells (*<u>CLU</u>, GFAP, <u>TMSB4X</u>, <u>CLDN4</u>, <u>TMSB10</u>, <u>CYR61</u>, <u>VIM</u>, <u>ELF3</u>, <u>IFITM3</u>, S100A10, <u>KRT8</u>, <u>S100A6</u>,* Figure 2D-E), all were previously reported in mouse inner ear tissues but only 7 in human inner ear tissues (underlined). To validate the specificity of these markers, we performed *in situ* hybridization and immunolabeled utricles procured from other organ donors and vestibular schwannoma subjects. Both anti-MYO7A and *SYT14* labeled all hair cells in organ donor and vestibular schwannoma samples (Figure 2F-G), where fewer hair cells and a flat epithelium were observed. As in mice, human supporting cell nuclei reside in the basal layer of the sensory epithelium and project cytoplasmic processes to the luminal surface (Figure 2F). As computationally predicted, both anti-GFAP and *ANXA2* marked only supporting cells from organ donor and vestibular schwannoma tissues (Figure 2F-G). Expression plots confirmed *ANXA2* and *SYT14* as pan-markers of supporting cells and hair cells, respectively (Figure 2H). Similarly, *in situ* hybridization validated *PPP1R27* and *LRRC10B* to be expressed in all hair cells and *CD9* in supporting cells (Figure S2E). These results indicate that gene expression in hair cells and supporting cells is well conserved between human organ donor and vestibular schwannoma utricles and between mice and humans.

**Figure 2.**
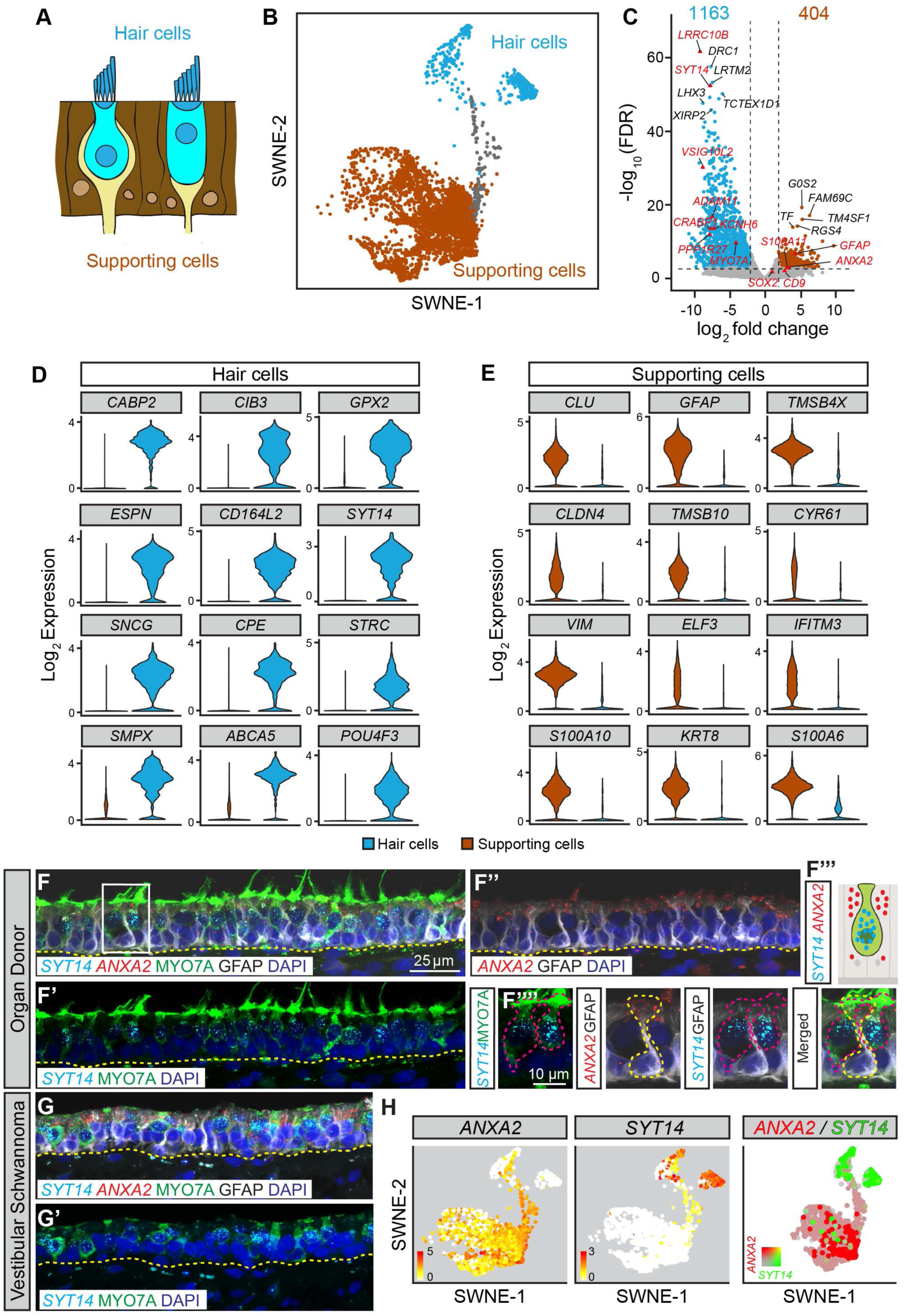
Differential gene expression in human vestibular hair cells and supporting cells. (A) Schematic of utricle sensory epithelium. (B) SWNE plot colored by hair cell and supporting cell groups with the hair cell precursor group (gray) excluded. (C) Volcano plot with enriched hair cell versus supporting cell genes using pseudobulk DESeq2 analysis. The top 5 differentially expressed genes are labeled as well as validated markers (red triangles). Using a log2 threshold of 2 and FDR < 0.01, 404 genes were enriched in the supporting cells and 1,163 in the hair cells (listed in Table S4). (D) Violin plots depicting 12 highly enriched hair cell (D) and supporting cell genes (E) using the Wilcoxon rank sum test. (F-G’) Validation of the hair cell marker *SYT14* (cyan) and supporting cell marker *ANXA2* (red) in utricles from organ donor and vestibular schwannoma patients. Cryosections were processed for fluorescent *in situ* hybridization and immunostaining for MYO7A (green), GFAP (gray) and DAPI (blue). Yellow dashed lines mark the basement membrane of the sensory epithelium. *SYT14* is expressed in MYO7A^+^ hair cells in (F’) and *ANXA2* is expressed in the cytoplasm of GFAP^+^ supporting cells in (F’’) in OD. (F’’’) Cartoon depicting the expression pattern of *SYT14* (perinuclear cytoplasm) and *ANXA2* (apical cytoplasm) in hair cells and supporting cells, respectively. Boxed area is magnified in individual panels in (F’’’’). Maroon dashed lines outline the MYO7A^+^/GFAP^−^ hair cells expressing *SYT14*. Yellow dashed lines outline the MYO7A^−^/GFAP^+^ supporting cells expressing *ANXA2*. (G-G’) *SYT14* marks hair cells and *ANXA2* supporting cells in vestibular schwannoma utricle, which is notably disorganized with loss of hair cells. (H) SWNE plots displaying enrichment of *ANXA2* in supporting cells and *SYT14* in hair cells. Log_2_ expression is shown with 0 in white and maximum in red at the indicated thresholds. Merged color SWNE plot demonstrates minimal overlap between these marker genes.

### Spatial and transcriptomic properties of hair cell subtypes

Utricular hair cells of both mouse and human fall into two main subtypes (~50% each of type I and II) (Rusch et al., 1998). In mice, this distinction is based on morphology, innervation, composition of ion channels, and gene expression (Burns et al., 2015; Jan et al., 2021; McInturff et al., 2018; Rusch et al., 1998). However, in humans, this configuration has been shown solely using morphological assessments of postmortem tissues (Hizli et al., 2016; Kaya et al., 2017). In our integrated dataset, differential gene expression revealed two putative hair cell subpopulations (Figure S2D), which we presumed to represent type I and type II hair cells (Figure 3B-C). Forty-seven genes were highly expressed human type I hair cells and 111 genes for type II hair cells (Figure 3C, Table S5, LFC>2 and FDR<0.01). Among the 13 most enriched type I hair cell genes (*VSIG10L2, BMP2, CLDN5, ALDH1A3, TAC1, FBXW7, CRABP1, ESPN, ATP2B2*, *PGA5, CHGA, PRKCD, ADAM11,* Figure 3D), ten have been reported in the mouse inner ear but only four showed expression restricted to type I hair cells (Burns et al., 2015; Jan et al., 2021). Reciprocally, we examined enriched genes in mouse utricular hair cell subtypes in our human dataset. The mouse type I hair cell marker *SPP1* (osteopontin) (McInturff et al., 2018), was notably not significantly enriched in our human dataset (Table S5). Except for *SOX2* and *BRIP1,* none of the enriched human type II hair cell genes (*CXCL14, CALB1, CSRP2, HTRA1, MORC1, SCNN1A, GNAS, NDUFA4L2, B3GNT10, CHRNA1,* Figure 3E) (Jan et al., 2021) were previously reported to be enriched in mouse type II hair cells in mice. Similarly, the established mouse type II hair cell markers *ANXA4* and *MAPT* (Burns et al., 2015; McInturff et al., 2018) were not enriched in the human type II hair cell cluster (Table S5). Unexpectedly, the mouse type I hair cell marker, *CALB1* (Stone et al., 2021), was enriched in human type II hair cells. These data suggest that the transcriptome profiles of hair cell subtypes diverge between mice and humans.

**Figure 3.**
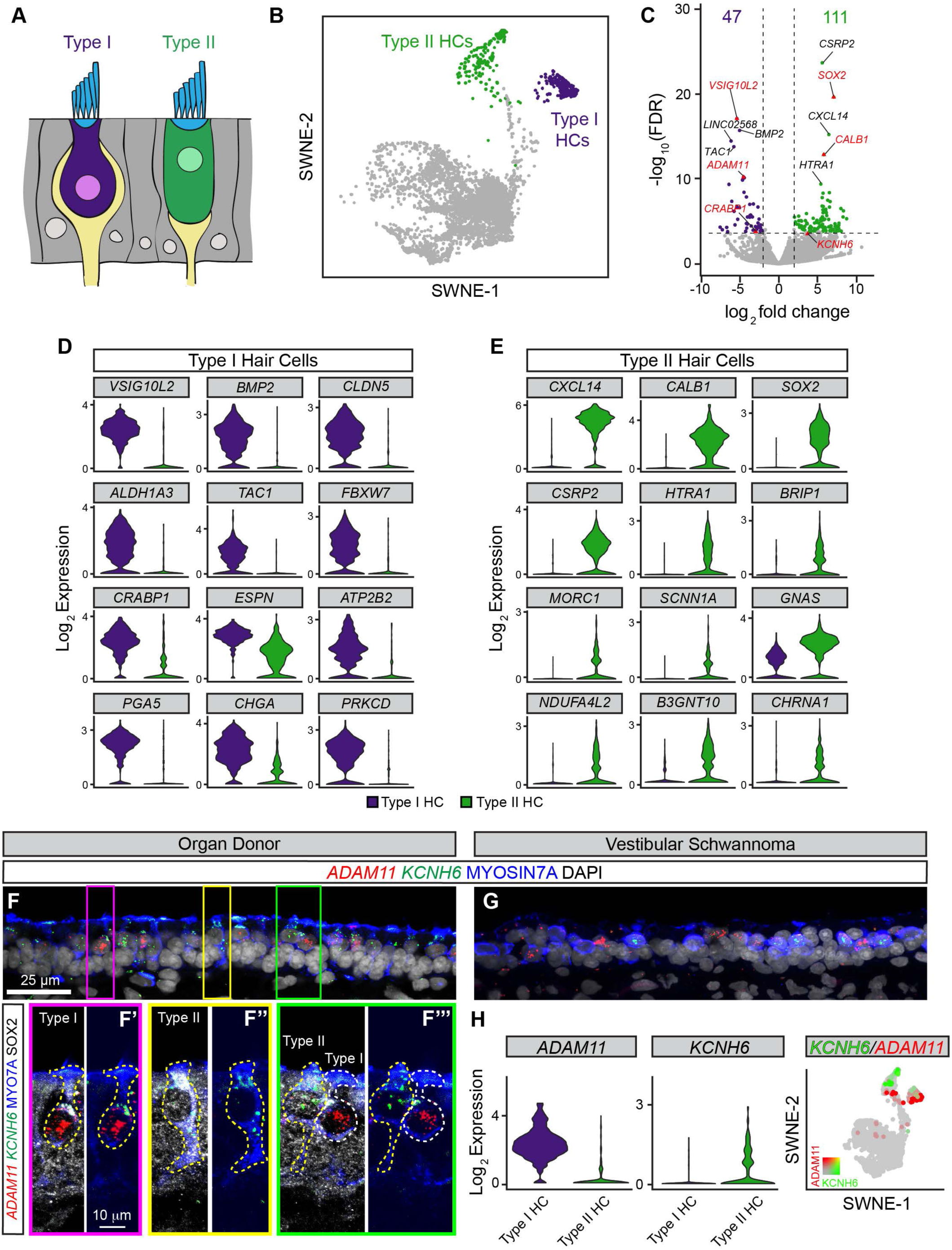
Human vestibular hair cell subtypes in vestibular schwannoma and organ donor utricles. (A) Schematic showing amphora-shaped type I hair cells with apical neck and calyceal innervation. Type II hair cells are goblet-shaped and display bouton-type innervation. (B) SWNE plot with distinct clusters of type I and II hair cells highlighted. (C) Volcano plots of 47 and 111 enriched genes in type I and II vestibular hair cells, respectively using DESeq2 pseudobulk analysis. The top 5 differentially expressed genes are labeled and validated genes are marked with red triangles. See Table S5 for complete list of genes. (D-E) Violin plots of enriched markers of type I and II hair cells using Wilcoxon rank sum test. (F-G) Combined fluorescent *in situ* hybridization and immunostaining validating expression of *ADAM11* in type I hair cells (red) and *KCNH6* in type II hair cells in sections of organ donor and vestibular schwannoma utricles counterstained with MYO7A (blue) and DAPI (gray). (F’) High magnification images of a type I hair cell from F (maroon box). This SOX2^−^ (gray) type I hair cell is amphora-shaped, displays a narrow apical neck, and robustly expresses *ADAM11*. (F’’) shows high magnification image of a type II hair cell from F (yellow box). This SOX2^+^ type II hair cell is goblet-shaped, displays basolateral process, and expresses *KCNH6.* (F’’’) Additional examples of type I and type II hair cells with *ADAM11* and *KCNH6* expression, respectively, from F (green box). (G) Type I and II hair cells marked by *ADAM11* and *KCNH6* in a vestibular schwannoma utricle, which has notably fewer hair cells of both types. (H) Violin and SWNE plots showing enrichment of *ADAM11* and *KCNH6* in type I and II hair cells, respectively.

We next compared the human utricular sensory epithelia of organ donors and vestibular schwannoma patients. In the mouse and human utricular sensory epithelium, hair cell subtypes have distinct shapes: type I hair cells are amphora-shaped while type II hair cells are goblet-shaped and display basolateral cytoplasmic processes. Both hair cell subtypes have their nuclei closer to the epithelial surface contrasting those of supporting cells that are closer to the basal surface (Hizli et al., 2016; Merchant et al., 2000; Pujol et al., 2014; Wang et al., 2019) (Figure 3A).

In organ donor tissues, the morphology of MYO7A^+^ hair cells was maintained and expression of *ADAM11* was exclusive to amphora-shaped type I hair cells. Type II hair cells displayed basolateral processes and were specifically marked by both anti-SOX2 and *KCNH6* (Figure 3F). In vestibular schwannoma patients, despite the loss of lamination and cell loss, both *ADAM11* and *KCNH6* expression are non-overlapping, and presumed restricted to type I and II hair cells, respectively (Figure 3G). These results validate the transcriptomic data showing enrichment of *ADAM11* and *KCNH6* in type I and II hair cells, respectively, in both cohorts (Figure 3H). Moreover, we validated expression of enriched type I hair cell (*CRABP1* and *VSIG10L2*) and type II hair cell genes (*CALB1*) (Figure S3A-B). These results confirmed the presence of two transcriptionally distinct human utricular hair cell subtypes in both organ donors and vestibular schwannoma utricles. Despite the conservation of morphology, human hair cell subtypes demonstrate divergent molecular features from mice.

### Diversity of supporting cell subtypes

Differential expression results have revealed several pan-supporting cells markers (Figure 2C, E), of which we have validated *GFAP* and *ANXA2* (Figure 2F). The mouse and human utricles are organized spatially into the central striolar and peripheral extrastriolar zones (Figure S3C) (Watanuki and Schuknecht, 1976) with distinct gene expressions described in mice (Jan et al., 2021; Ono et al., 2020). The striolar region may be more critical for utricular function as mutant mice missing this domain lack vestibular responses (Ono et al., 2020). Differential gene expression shows 17 genes enriched in striolar supporting cells and 37 genes in extrastriolar supporting cells (Figure S3E-F). *TECTB*, which marks striolar supporting cells in mouse and chicken utricles (Burns et al., 2015), was also enriched in human striolar supporting cells (Table S6). Only a small set of enriched genes marking human extrastriolar (two of 12, *ALDH1A1, S100B)* and striolar (six of 12, *SFRP2*, *TECTB, ISL1, RGCC, CA8 and GATA3*) supporting cells were previously reported in the mouse utricle (Burns et al., 2015; Jan et al., 2021). Therefore, like human hair cell subtypes, human supporting cell subtypes also display divergence from those in mice.

### Hair cell degeneration in human vestibular schwannoma utricle

Postmortem studies on utricles from vestibular schwannoma patients have reported hair cell degeneration (Hizli et al., 2016; Sans et al., 1996). To independently verify these results and determine whether supporting cells mount a regenerative response, we first immunolabeled utricles for hair cells (MYO7A^+^) and supporting cells (MYO7A^−^ and SOX2^+^) from organ donor (n=5, 4 males and 1 female, aged 42.0±25.5) and vestibular schwannoma subjects (n=13, 9 males and 4 females, aged 50.5±16.7). As a secondary control, we included utricles from cadavers (n=4, 2 males and 2 females, aged 75.8±5.9, Table S1) without any history of auditory and vestibular deficits. Compared to organ donors or cadavers, vestibular schwannoma utricles contained significantly fewer hair cells (~78% fewer, p<0.001, Figure 4A-D, S4A-B’’, D-G, Table S7) and supporting cells (~21% fewer, p<0.05, Figure S4A’-C, Table S7). Hair cell counts from individual vestibular schwannoma patients were comparable despite variability of age, gender, laterality, tumor size, or history of radiation in the cohort (Figure S4F, Table S1). Collectively, these results further indicate that vestibular schwannoma utricles represent diseased organs whereas organ donor and cadaveric organs serve as normal controls.

**Figure 4.**
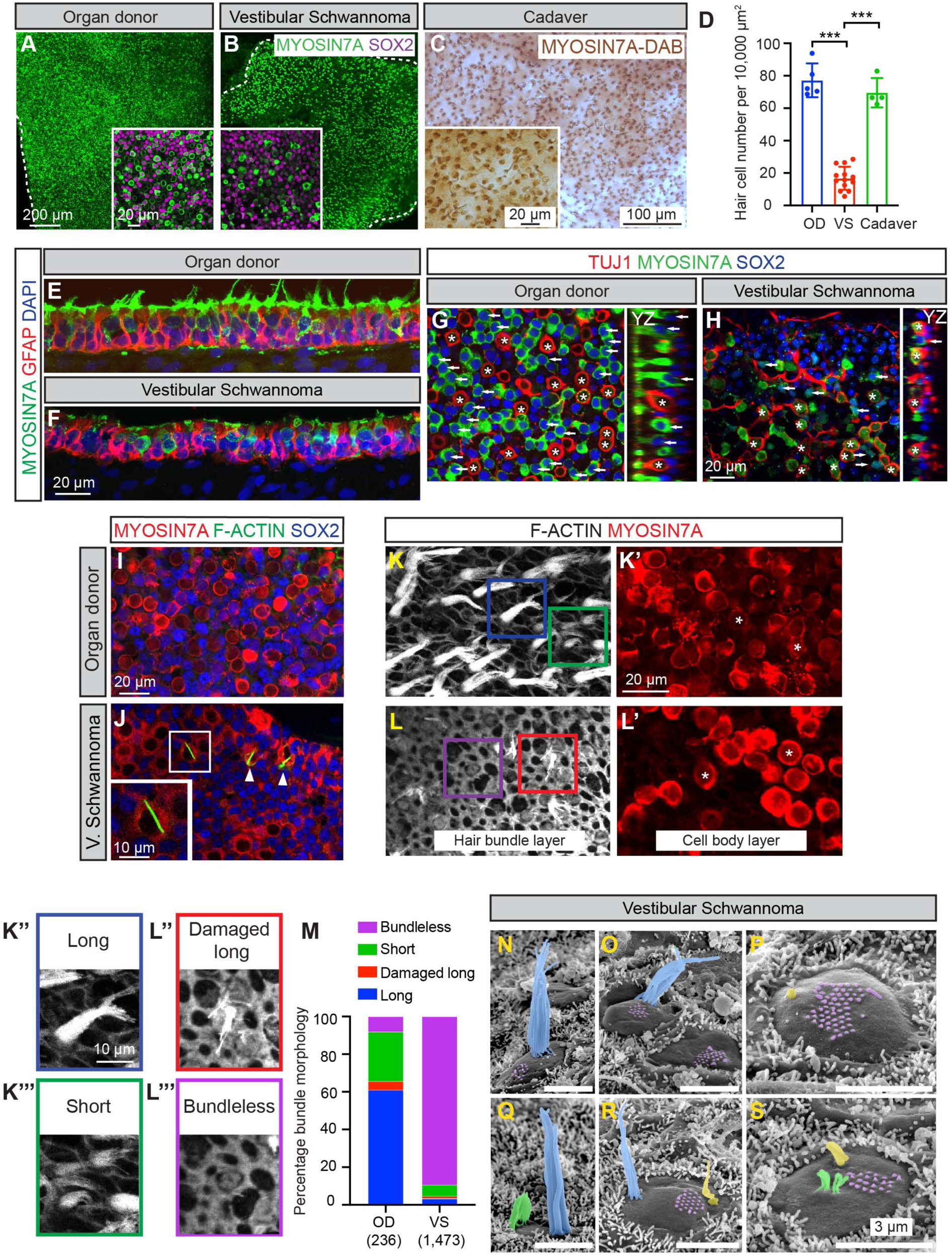
Hair cell degeneration in human vestibular schwannoma utricles. (A-C) Whole mount preparation of utricles from organ donors and vestibular schwannoma patients. Representative low magnification images of MYO7A^+^ hair cells (green) in the sensory epithelium from both groups (dashed lines). Insets show representative high magnification images of hair cells and supporting cells (magenta). Fewer hair cells were observed in vestibular schwannoma than organ donor tissues. (C) Whole mount preparation of utricular sensory epithelium from cadavers. MYO7A-DAB (brown) staining showing a dense population of hair cells. Inset shows a high magnification image. (D) Quantification showing that vestibular schwannoma utricles (n=13) had significantly fewer hair cells than either those from organ donors (n=5) or cadavers (n=4). (E-F) Representative images of sections of utricular sensory epithelium of organ donor and vestibular schwannoma patients. Compared to organ donor utricles, the vestibular schwannoma sensory epithelium appeared thinner, contained fewer hair cells (MYO7A, green), with many supporting cells (GFAP, red) remaining. Hair cells in the vestibular schwannoma sensory epithelium appeared dysmorphic and devoid of bundles. (G-H) Relative to organ donor utricles, vestibular schwannoma utricles displayed fewer type I hair cells (asterisks, SOX2^−^ (blue), TUJ1^+^ calyces) and type II hair cells (arrow, SOX2^+^). (I-J) F-actin (green) labeling reveals degenerating hair cells (arrowhead and inset, cytocauds with actin-rich cables) in VS but not OD tissues. (K-L’) Representative images showing hair cells with damaged bundles (F-actin, gray) connected to MYO7A^+^ cell bodies (red, asterisks) in VS but not OD tissues. (K’’-L’’’) Representative high magnification images showing 4 types of bundle morphology observed in K and L: long bundles (blue box), damaged long bundles (red box), short bundles (green box) and bundle-less (purple box). (M) Percentage of hair cells displaying distinct bundle morphology in hair cells from OD and VS tissues. Most hair cells are bundle-less in VS tissues (n=1,473 cells from 3 samples), whereas many hair cells in OD tissues have long bundles (n=236 from 1 sample). (N-O) Scanning electron microscopy images of hair cells from a VS utricle. (N) Hair cell with normal-appearing hair bundle (colored in blue). (O) Two hair cells with remnants of hair bundles visible as short stereocilia stubs at the apical surface (colored in purple); the left hair cell has an additional normal appearing hair bundle. (P) Remnants of stereocilia and kinocilium (colored in yellow). (Q) Hair cell with short (green) and tall hair bundles located at the opposite poles of the apical surface. (R) Hair cell with few normal appearing stereocilia and remnants of a hair bundle at the opposite apical pole. (S) Hair cell with a few thin and short stereocilia and remnants of a hair bundle. Data shown as mean±S.D. and compared using one-way ANOVA. ***p<0.001.

To characterize the degree of degeneration, we immunolabeled sections of organ donor and vestibular schwannoma utricles for the hair cell marker MYO7A and supporting cell marker GFAP. Compared to organ donor utricles, vestibular schwannoma utricles contained fewer hair cells and appeared thinner, resulting in less lamination of hair cell and supporting cell nuclei (Figure 4E-F). Hair cells in the vestibular schwannoma sensory epithelium appeared short and round, and many lacked hair bundles (Figure 4E-F). Using TUJ1^+^ calyces or SOX2 protein expression as markers of type I and II hair cells, respectively, we found that vestibular schwannoma organs had significantly fewer of both type I and II hair cell subtypes than organ donor tissues. However, there was disproportionally more type I hair cells in the vestibular schwannoma than organ donor utricles (3.51 versus 0.99, respectively, Figure 4G-H, S4H-I, Table S7), suggesting a previously unrecognized, preferential loss of type II hair cells.

We observed cytocauds, which are actin-rich cables in degenerating hair cells (Kanzaki et al., 2002; Taylor et al., 2015), in vestibular schwannoma but not organ donor utricles, suggesting ongoing hair cell degeneration in the former (Figure 4I-J, S4J-J’). While phalloidin-labeled bundles were apparent in most MYO7A^+^ hair cells in the organ donor utricle, few bundle-bearing hair cells were observed in the vestibular schwannoma utricles (10.7% of 1,473 hair cells from 3 vestibular schwannoma utricles, Figure 4K-M). Remnant hair bundles appeared damaged (splayed) or short (13.4% and 56.1% of 157 hair cells from 3 vestibular schwannoma utricles, Figure 4M). This contrasts with the organ donor utricles where 89.3% of hair cells displayed bundles, with most appearing long (61.1%, n=236 hair cells from 1 utricle, Figure 4K-M, S4K-L). Under scanning electron microscopy, hair bundles with a staircase pattern and packed stereocilia were rarely found (Figure 4N) in vestibular schwannoma utricles, whereas others lacked stereocilia and kinocilium and instead displayed remnants of short bundles at the apical surfaces (Figure 4O-P, S4M-O). In other cases, stereocilia within a hair bundle had fused (Figure 4Q). Lastly, we found individual hair cells with two distinct hair bundles located at the opposite poles of the apical surface (Figure 4Q-S), one with taller and thicker stereocilia than the other, possibly corresponding to different maturation stages, perhaps an indication of ongoing repair or regeneration that has been previously suggested (Kanzaki et al., 2002; Taylor et al., 2015). Together, these results indicate ongoing hair cell degeneration and hair bundle damage in the vestibular schwannoma utricles.

### Transcriptomic and trajectory analysis of hair cell regeneration in the human utricle

Both the mouse and human utricle demonstrate a limited degree of hair cell regeneration by supporting cells (Forge et al., 1993; Golub et al., 2012; Sayyid et al., 2019; Taylor et al., 2018; Warchol et al., 1993), although spontaneous regeneration in vestibular schwannoma utricle has not been directly observed. To define the molecular underpinning of human hair cell regeneration, we performed dimension reduction using SWNE on the integrated dataset. The SWNE plot allowed visualization of presumed hair cell precursors (cluster 20, Figure 5A) that displayed spectra of hair cell and supporting cell marker genes. To delineate genes directing supporting cell to hair cell transition, we created unbiased trajectories (Figure S5B), focusing on two lineages that originate from supporting cells to type I and type II hair cells via the intermediate hair cell precursors (Figure 5A, S5B).

**Figure 5.**
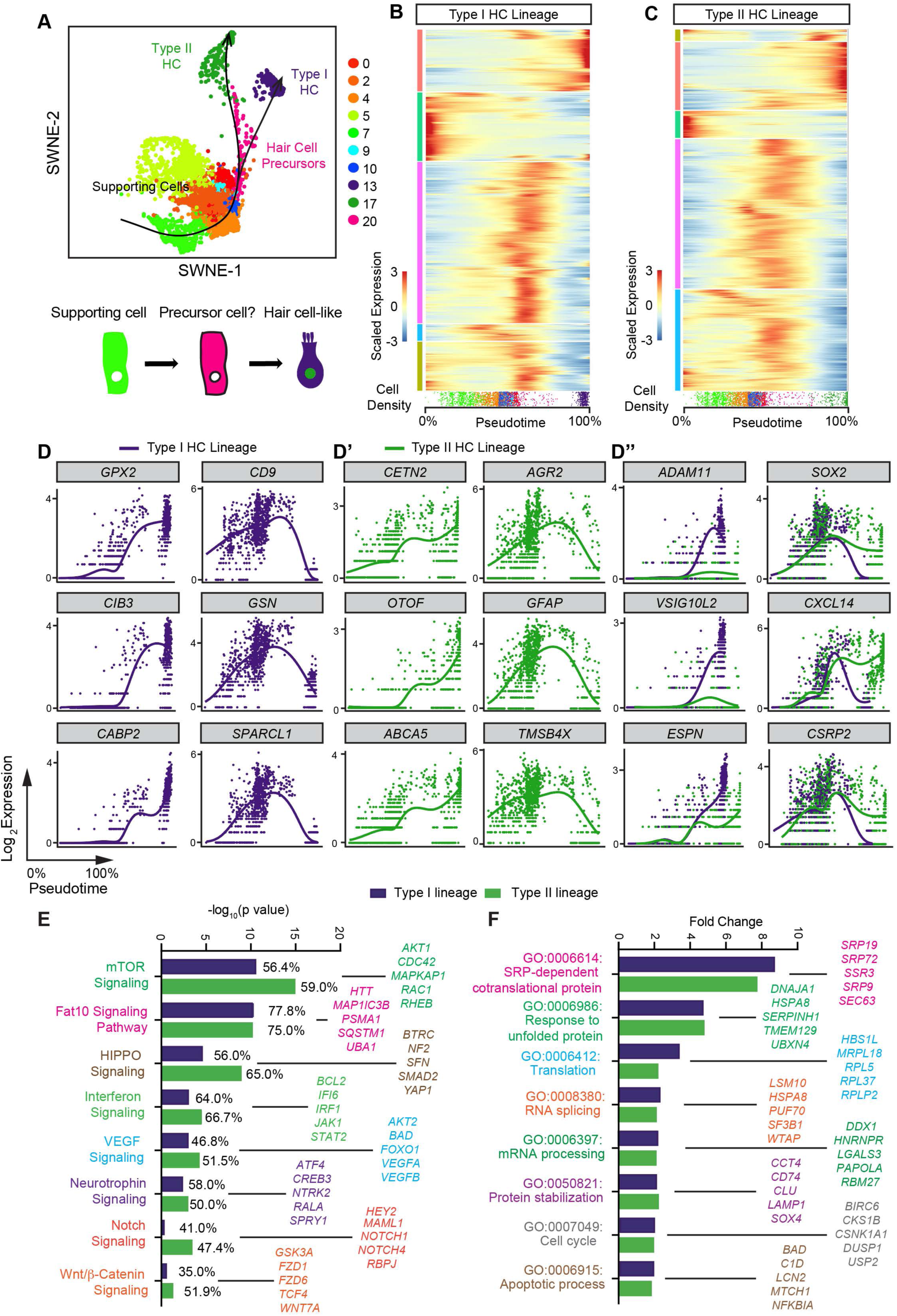
Differential gene expression along trajectories linking supporting cells and hair cells in organ donor and vestibular schwannoma utricles. (A) SWNE plot with projected type I and type II hair cell lineages using *Slingshot* starting at the supporting cell clusters. Predicted hair cell precursors are labeled in pink, cluster 20. (B-C) Heatmap depicting genes that are dynamically changing along the type I and II hair cell lineages. An association test in *tradeSeq* was used to determine statistically significant genes changing along pseudotime with an FDR ≤ 0.01, revealing 2,787 and 7,445 genes in type I and type II hair cells, respectively. There are five patterns of dynamic expression with some genes upregulated, some downregulated, and others transiently expressed in both type I and II lineages. The density of cells and corresponding location along the lineage is depicted below the heatmap. Genes listed in Tables S8-9. (D-D’) Plots showing the dynamic expressions of genes from (B-C) that increase (left column) or decrease (right column) along pseudotime as a supporting cell transitions to a type I or II hair cell. The individual dots in each plot represent single cells with the x-axis representing pseudotime and the y-axis showing log_2_ expressions. The solid line represents the generalized additive model fit to the expression pattern for each gene. (D”) Comparison of type I versus type II hair cell dynamic gene expressions. Using the different end test, we detected statistically significant genes that begin at similar expression levels and end at different levels. *SOX2*, *CXCL14*, and *CSRP2* expression increase in type II hair cells, while *ADAM11*, *VSIG10L2*, and *ESPN* increase in type I hair cells. (E) Graph showing eight of the top scoring canonical pathways associated with the type I and/or type II lineages along with the corresponding percentage of significantly expressed genes relative to the total number of genes in each pathway. (F) Graph of top biological function GO terms associated with a type I and type II lineages.

Using a generalized additive model fit onto the two lineages of type I and type II hair cells, we identified 2,787 and 7,445 significant, dynamically expressed genes along the two respective trajectories (Figure 5B-C). An association test detected five patterns of dynamic gene expression in type I and II hair cell lineages (Figure 5B-C, Table S8-9): rapidly upregulated, rapidly downregulated, gradually upregulated, gradually downregulated, and transiently upregulated (Figure 5B-C). Both the type I and II lineages display rapidly upregulated and downregulated genes (type I: *GPX2*, *CIB3*, *GABP2* and *CD9*, *GSN*, *SPARCL1*; type II: *CETN2*, *OTOF*, *ABCA5* and *AGR2*, *GFAP*, *TMSB4X*, Figure 5D-D’), implicating shared features of hair cell specification and differentiation. To identify divergent genes among the type I and type II lineages, we computed the Different End Test and identified genes upregulated solely in type I hair cells (*ADAM11*, *VSIG10L2*, and *ESPN*) and type II hair cells (*SOX2*, *CXCL14*, and *CSRP2*, Figure 5D’’). These results suggest that regeneration of type I and II hair cells share some but not all molecular features.

To characterize mechanisms driving supporting cell-to-hair cell transition, we analyzed dynamically expressed genes in type I and II lineages and found 196 and 297 associated canonical pathways, respectively (Fisher’s Exact Test, p<0.05, Table S10-11), most of which were shared by both lineages. Shared pathways included FAT10, mTOR, VEGF, Neurotrophin, Interferon, HIPPO signaling, while Notch and Wnt/ß-catenin signaling were only associated with the type II lineage (Figure 5E). In mice, HIPPO, Notch and Wnt signaling have been shown to regulate regeneration of the utricle, but their effects on specification of type I versus type II hair cells are not known (Lin et al., 2011; Rudolf et al., 2020; Wang et al., 2015). Moreover, 24 and 15 biological function GO terms were identified for dynamically expressed genes in the type I and II lineages (FDR<0.05, Figure S5F, Table S12-13). These results serve as a mechanistic framework for characterizing naturally occurring human hair cell regeneration.

### Vestibular schwannoma utricles contain more hair cell precursors than organ donor utricles

To characterize hair cell precursors, we examined dynamically expressed genes as supporting cells transition into hair cells in both the type I and II lineages. As markers of both type I and II hair cells (Figure 2), *SYT14* and *MYO7A* were upregulated along both lineages (Figure 6A). Conversely, the supporting cell marker *ANXA2* (Figure 2) was downregulated along these lineages (Figure 6A). In vestibular schwannoma utricles, where almost all hair cells expressed *SYT14* and *MYO7A* and supporting cells expressed *ANXA2*, we found occasional elongated cells co-expressing *ANXA2*, *SYT14*, and MYO7A (Figure 6B-B’’’’), suggesting a transitory cell state between a supporting cell and hair cell *in vivo*.

**Figure 6.**
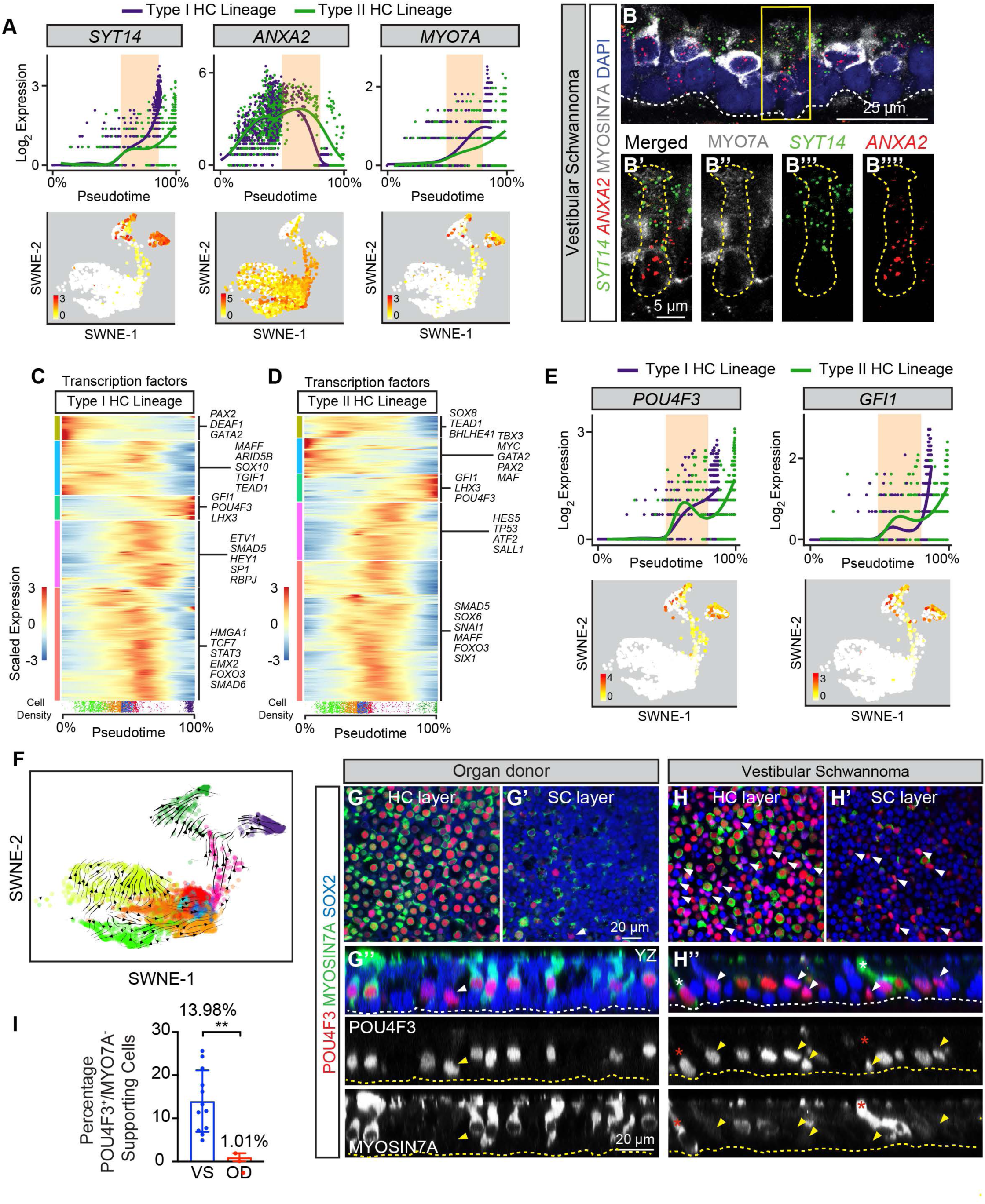
Regenerative responses in organ donor and vestibular schwannoma utricles. (A) Pseudotime plots with generalized additive models depicting dynamic gene expressions along the supporting cell-to-type I hair cell and -type II hair cell lineages. The location of hair cell precursors along pseudotime is highlighted (peach). These plots predict that hair cell precursors co-express *SYT14*, *ANXA2*, and *MYO7A*. Both *SYT14* and *MYO7A* expression is upregulated while *ANXA2* expression is downregulated in both lineages. SWNE expression plots show the 2-dimensional locations of cells expressing these genes. (B-B’’’’) Combined *in situ* hybridization and immunostaining of cryosection of a vestibular schwannoma utricle, showing an elongated hair cell-precursor-like cell expressing *SYT14* (green), *ANXA2* (red), MYO7A (gray). White dashed line marks the basement membrane of the sensory epithelium. Boxed area is magnified in individual panels in (B’-B’’’’). A *SYT14*, *ANXA2* and MYO7A triple positive cell (yellow dashed line) was observed. The nucleus of this cell is located close to the basement membrane, below that of other hair cells. (C-D) Dynamic expression of transcription factors along the type I and II hair cell lineages. The association test revealed 446 transcription factors with statistically significant dynamic expression patterns in each of type I and type II trajectories. Sample genes from each of 5 patterns of expression are shown. The location of cells along pseudotime for each lineage is depicted as cell density below the heatmaps. (E) Dynamic expression and SWNE plots predicting upregulation of both *POU4F3* and *GFI1* as supporting cells transition to both type I and II hair cells. (F) RNA velocity analysis shows the hair cell precursor group is differentiating towards type I and type II hair cells and away from supporting cells. (G-H’’) Whole mount preparation of utricles from both OD and VS patients were immunostained for POU4F3 (red), MYO7A (green) and SOX2 (blue). Many POU4F3^+^/SOX2^+^/MYO7A^−^ hair cell-precursor cells (arrowheads) were present in the VS tissues, but only few were detected in OD tissues. Representative orthogonal views show POU4F3^+^/SOX2^+^/MYO7A^−^ cells (arrowhead) with nuclei near the basement membrane. Occasional elongated POU4F3^+^/SOX2^+^/MYO7A^−^ cells (asterisks) were found in VS utricle (H’’). These hair cell-precursor cells display nuclei at the supporting cell level and below that of other hair cells. (I) The percentage of POU4F3^+^/MYO7A^−^ supporting cells was significantly higher in VS (n=12) than to OD (n=3) tissues. Data shown as mean ± S.D. and compared using Student’s t-tests. **p<0.01.

To confirm the presence of hair cell precursors *in vivo* and to further identify drivers of hair cell regeneration, we examined transcription factors among the identified dynamically expressed genes (Hu et al., 2019). A total of 446 significant dynamically expressed transcription factors were identified in both type I and II lineages (Figure 6C-D, Table S14-15). These include *POU4F3* and *GFI1* which are required for hair cell survival and maturation in mice (Erkman et al., 1996; Hertzano et al., 2004) (Figure 6C-D). Surprisingly, *ATOH1*, which promotes hair cell formation in mouse and human utricles (Sayyid et al., 2019; Taylor et al., 2018), was found modestly expressed in a subset of hair cells and not dynamically expressed (Figure S6K-K’, L). Because *POU4F3* and *GFI1* were rapidly upregulated along the type I and type II hair cell trajectories (Figure 6E), we postulated that they would label both hair cell precursors and differentiated hair cells. As an independent computational method of validating supporting cell-hair cell transition, we performed RNA velocity analysis (Bergen et al., 2020; La Manno et al., 2018) on our subsetted data. In this method, each individual cell’s trajectory is computed irrespective of surrounding cells’ trajectories using the ratio of spliced to unspliced mRNA to derive the RNA velocity on a gene-by-gene level. Our data showed that the hair cell precursor group is projected to differentiate from a supporting cell state towards the type I and type II hair cell states (Figure 6F, S6M), further implicating that supporting cell-hair cell transition occurs *in situ*. As expected, in organ donor and vestibular schwannoma utricles, all MYO7A^+^ hair cells were marked with anti-POU4F3 and anti-GFI1 antibodies. Strikingly, a subset of SOX2^+^, MYO7A^−^ supporting cells expressed POU4F3 or GFI1 protein and mRNA in both the organ donor and vestibular schwannoma utricles (Figure 6G-H, S6A-G’, I-J), with the latter displaying significantly more POU4F3 (13.98 vs. 1.01%, p<0.01, Figure 6I) or GFI1-positive hair cell precursors (12.79 vs. 0.19%, p<0.05, S6H). Collectively, these results unveil transcriptomes of human hair cell precursors, which were more prevalent in vestibular schwannoma than organ donor utricles *in situ*.

## Discussion

Inner ear sensory hair cells are specialized mechanoreceptors required for hearing and balance functions. Unlike the mammalian cochlea, the vestibular organs have a limited capacity to regenerate lost hair cells (Atkinson et al., 2015). Rodents have served as the primary model system examining mammalian vestibular hair cell regeneration, relying on toxins or transgenic approaches to induce damage (Golub et al., 2012; Sayyid et al., 2019). Prior studies using tissues procured from vestibular schwannoma patients showed that supporting cells can spontaneously regenerate hair cells after gentamicin treatment; likewise, ectopic hair cells can form in response to ATOH1 overexpression (Taylor et al., 2018; Taylor et al., 2015; Warchol et al., 1993). Here, we procured utricles from organ donor and vestibular schwannoma subjects. The former cohort lacked any history of auditory or vestibular dysfunction, while the latter exhibited hearing loss and/or vestibular dysfunction. Single-cell transcriptomic analysis and *in vivo* validation experiments revealed conserved transcriptomes of hair cells and supporting cells, despite significant and ongoing degeneration in the vestibular schwannoma organs. Despite conservation of generic hair cell and supporting cell genes between mice and humans, we found significant interspecies divergence among hair cell and supporting cell subtypes. By analyzing dynamically expressed genes in the supporting cell-hair cell trajectories, we predicted *POU4F3* and *GFI1* to mark hair cell precursors. We validated these markers and discovered that significantly more hair cell precursors reside in the vestibular schwannoma than organ donor utricles. Our study reveals candidate mechanisms driving spontaneous human hair cell regeneration and serves as a data-rich resource for research on human auditory and vestibular disorders.

### Single cell transcriptomic studies on the inner ear

Single-cell transcriptomic analysis has been instrumental in defining cellular heterogeneity in the mouse inner ear as well as discovering candidate mechanisms of development, hair cell regeneration and pathogenesis of auditory dysfunction (Wu et al., 2021). While these studies can serve as foundational models for human disease, studies on the human inner ear have almost exclusively relied on postmortem, cadaveric tissues (Hizli et al., 2016; Kaya et al., 2017; Lopez et al., 2016a; Merchant, 1999; Merchant et al., 2000; Sans et al., 1996; Senn et al., 2020; Watanuki and Schuknecht, 1976) or tissues originating from fetuses or diseased inner ears. In the current study, transcriptomes of distinct cell types from the healthy and diseased mature, human inner ear serve as benchmarks guiding future works.

Because the mouse and human inner ear organs share histologic features and constituents essential for hearing and balance functions (Avraham, 2003; Kikkawa et al., 2012; Ohlemiller, 2019), mouse models are commonly used to study human auditory and vestibular diseases. Several pathways posited to regulate human vestibular hair cell regeneration (Wnt/ß-catenin, Notch, HIPPO) were previously linked to regeneration in the mouse vestibular system (Lin et al., 2011; Rudolf et al., 2020; Wang et al., 2015), while others have unknown functions (FAT10, mTOR, VEGF, Neurotrophin, Interferon) (Figure 5). Remarkably, our data suggest that regeneration of type I and II hair cells may be differentially regulated by Notch and Wnt/ß-catenin signaling and warrant further testing. When comparing enriched genes between human and mouse utricles (Burns et al., 2015; Jan et al., 2021; McInturff et al., 2018), we found overlap among generic hair cells and supporting cells but striking divergence among hair cell and supporting cell subtypes. Such an interspecies difference that has previously been suggested in the refined cell subtypes of the mouse and human immune systems (Cheng et al., 2014; Gilbertson and Weinmann, 2021). Along a similar vein, divergence of neurotransmitter receptors and ion channels is known between mouse and human brains (Hodge et al., 2019). Such a divergence has implications for translating mouse studies to humans and emphasizes the importance of investigating human auditory and vestibular organs.

### Application of human utricle single-cell data to study hearing loss and vestibular diseases

Our study serves as an invaluable resource to study human inner ear diseases. As a case in point, we interrogated the expression patterns of 70 human hearing loss- and 20 vestibulopathy-related genes in our integrated dataset, 10 of which were linked to both conditions. By determining the mean relative expression of these genes, we found that 40 hearing loss genes are uniquely expressed in hair cells and that seven non-sensory cell types highly expressed 30 hearing loss genes (Figure S7A). Among the 20 vestibulopathy-related genes, seven are uniquely expressed in hair cells and 13 in seven non-sensory cell types (Figure S7B). In contrast to the rich literature on the pathogenesis of hearing loss genes in hair cells, studies on non-sensory cell types and vestibulopathy-related genes are scarce, and our study establishes a foundation for these future studies.

### Regeneration in human vestibular schwannoma utricles

Inner ear tissues collected from vestibular schwannoma patients have served as the primary model system to study human hair cell regeneration (Taylor et al., 2018; Taylor et al., 2015; Warchol et al., 1993). Although hair cells are known to have degenerated in inner ear tissues from vestibular schwannoma patients (Hizli et al., 2016; Sans et al., 1996), these studies on regeneration have employed ototoxins to ablate remaining hair cells *in vitro*. By comparing vestibular schwannoma to organ donor utricles, we discovered that the former represents damaged tissues with active regeneration *in situ* and therefore is a distinct model system of human hair cell degeneration and regeneration.

In mice, lineage-tracing experiments show that regenerating hair cells are primarily type II and that they incompletely differentiate (Golub et al., 2012; Sayyid et al., 2019; Wang et al., 2019). Our results suggest that regenerating human hair cells display a spectrum of differentiation. While these results may indicate that human vestibular hair cell regeneration, as in mice, requires exogenous factors to enhance the degree of regeneration and differentiation, it is also possible that ongoing damage from vestibular schwannoma modulates these processes.

In summary, using vestibular tissues procured from adult human organ donors and vestibular schwannoma patients, we have revealed the transcriptome of inner ear sensory and non-sensory cells. We show that vestibular schwannoma utricles harbor both degenerating sensory cells and hair cell precursors. The adult human inner ear has to date remained an enigma at the molecular level, and it is of utmost public health importance to decipher mechanisms of damage and repair. Our results present a foundational resource for future studies on human auditory and vestibular function and diseases.

## Methods

### Human vestibular organ procurement

Whole organ utricles were collected intraoperatively from patients with vestibular schwannoma and organ donors. For vestibular schwannoma patients, utricles were procured from patients undergoing translabyrinthine resection of tumors. Briefly, a neurotologist (R.K.J., P.S.M., N.B.) opened the vestibule by drilling with a diamond burr on low speed, microdissected out the utricle and removing the attached ampullae. The utricle was then placed in phosphate buffered solution (PBS, pH 7.4; Electron Microscopy Services) on ice until further analysis (<10 min).

Organ donors were referred by Donor Network West (San Ramon, CA). Bilateral utricles were harvested from organ donors as previously described (Aaron et al., 2022; Vaisbuch et al., 2022). Briefly, a post-auricular incision was made followed by a transcanal approach to expose the middle ear. The tympanic membrane, malleus and incus were removed while keeping the stapes *in situ*. To expose the vestibular organs, bony covering of the vestibule was thinned using a diamond burr on low speed, the stapes footplate removed, followed by widening of the oval window. The utricle was harvested from the elliptical recess and placed in PBS on ice for single-cell RNA sequencing analysis or in 4% paraformaldehyde for histologic analysis.

For cadaveric utricles, temporal bones were collected 10-14 hours postmortem, following which specimens were microdissected from temporal bones acquired at autopsy as previously described (Lopez et al., 2005).

### Human subject inclusion criteria

Three cohorts were included in the study. In the first cohort, twenty-one patients (21 ears) with vestibular schwannoma were enrolled from Stanford University (Palo Alto, CA). Adult subjects with a vestibular schwannoma diagnosed by MRI and hearing loss documented by audiograms were included. Those undergoing a translabyrinthine approach for tumor resection between March 2015 to August 2021 were enrolled into the study. The second cohort were 8 ears from 6 (pediatric and adult) organ donors enrolled through Donor Network West (San Ramon, CA) between January 2018 and July 2020. Both cardiac and brain death donors with no known history of otologic disorders were included. The last cohort consisted of 4 ears from 4 postmortem adult cadavers from University of California, Los Angeles (Los Angeles, California) between February 2015 to January 2017.

All protocols were approved by the Institutional Review Board of the Stanford University (IRB 27500, 48579, 38993, 50076), Donor Network West’s internal ethics committee (IRB #STAN-17-200) and its medical advisory board, and University of California, Los Angeles (IRB 10-001449).

### Clinical information

The following was collected for all cohorts: age, gender, and laterality (Table S1). For the first cohort, we also collected information pertaining to vestibular symptoms (dizziness, vertigo, or imbalance) or physical signs (positive Romberg sign or unsteady gait), tumor size (longest diameter and its orthogonal measurement in the axial view of the cisternal component), pure tone averages (PTAs) (calculated using 0.5, 1, 2, and 4 kHz), word recognition scores, and history of radiation (Table S1).

### Cryosections

Tissues were fixed in 4% paraformaldehyde (PFA) (PBS pH 7.4; Electron Microscopy Services) for 16-24 hrs at 4°C. After washing in PBS, tissues were cryoprotected in a sucrose gradient (15%, 20% and 30% for 1 hour each), embedded in 100% OCT, and frozen on dry ice. Sections were cut at 10-15 μm and frozen at −80°C.

### Immunohistochemistry (Lopez et al., 2016b; Wang et al., 2019)

Utricles from vestibular schwannoma patients and organ donors were fixed in 4% PFA (in PBS, pH 7.4; Electron Microscopy Services) for 40 min at RT or 20 hr at 4°C. Tissues were blocked with 5% goat or donkey serum, 0.1% tritonX-100, 1% bovine serum albumin (BSA), and 0.02% sodium azide (NaN_3_) in PBS at pH 7.4 for 1-2 hrs at room temperature, followed by incubation with primary antibodies diluted in the same blocking solution overnight at 4°C. The next day, after washing with PBS, tissues were incubated with secondary antibodies diluted in 0.1% triton X-100, 0.1% BSA, and 0.02% NaN_3_ solution in PBS for 2 hr at room temperature. After PBS washing, specimens were mounted in antifade Fluorescence Mounting Medium (DAKO) or ProlongGold (Thermo Fisher Scientific) and coverslipped. We used antibodies against the following markers: MYOSIN7A (1:1000-5000; Proteus Bioscience), SOX2 (1:200-500, R&D and 1:400 Santa Cruz Biotechnology), POU4F3 (1:400, Santa Cruz Biotechnology), GFI1 (gift from H. Bellen), GFAP (1:250-1000, Sigma), TUJ1 (1:1000, Neuromics), and CALBINDIN1 (1:500, Cell Signaling Technologies). Secondary antibodies were conjugated with FITC (1:500, Thermo Fisher Scientific), TRITC (1:500, Thermo Fisher Scientific), CY5 (1:250, Thermo Fisher Scientific), Alexa 594 (1:500, Thermo Fisher Scientific) or Alexa 405 (1:250, Abcam). Fluorescent-conjugated phalloidin (1:1000; Sigma) and DAPI (1:10000; Invitrogen) were used.

Cadaveric temporal bones were fixed in 10% formalin for 6-24 months before microdissection. Utricles were then microdissected, placed individually in a rotary shaker, and incubated for 3 hours in a blocking solution containing 2% BSA fraction V (Sigma), 0.1% TritonX-100 (Sigma) diluted in PBS and incubated at 6°C. Subsequently, the blocking solution was removed and tissues were incubated with anti-MYOSIN7A in a rotatory shaker for 72 hrs at 4°C. Next, tissues were washed with PBS (20 min x 5) and incubated with the secondary biotinylated antibodies (1:1000 in PBS, Vector Labs) for 2 hrs at RT. Afterwards, tissues were washed again in PBS (20 min x 5) and incubated with the ABC complex (Vector Labs). The antigen-antibody reaction was visualized using 3,3′-Diaminobenzidine (DAB) (Vector Labs). Lastly, utricles were washed with PBS (20 min x 5) and flat mounted on glass slides and coverslipped with hard mount VECTASHIELD solution. Cryosections of mouse utricles subjected to the same protocol were used as positive controls. As negative controls, the primary antibody was omitted or preabsorbed with the antigen and the immunoreaction was performed as described above. No immunoreaction was detected in both types of negative controls.

### Image acquisition and analysis

Images of whole mounts and sections were acquired using confocal microscopy (LSM700 or LSM880, Carl Zeiss Microscopy), Axioplan 2 microscope coupled to a MRC5 (bright field) camera and using Axiovision AC software (Release 4.8, Carl Zeiss). Z-stack images were taken at 1 µm intervals. Data was analyzed with Image J64 (Fiji, NIH) and Adobe Photoshop (Creative Cloud, Adobe Systems).

### *In situ* hybridization

Detection of transcripts for *ATOH1*, *POU4F3* and *GFI1* was performed using the V2.5 HD Red *RNAscope™* detection system (322350, ACDBio, Newark, CA). The following probes from ACDBio were used: DAPB (negative control; Cat# 310043); POLR2A (positive control, Cat# 310451); ATOH1 (Cat# 417861); POU4F3 (Cat# 519131); GFI1 (Cat# 422981). Experiments were performed according to the manufacturer’s instructions, with the following modifications: slides were pre-baked at 60°C for 30 min before tissue processing, boiling was performed for 90 seconds, and the Protease Plus was diluted 1:3. In some cases, protease digestion was carried out at different temperatures to appropriately balance penetration of probes with retention of immunohistochemical antigens.

Detection of additional RNA transcripts was performed using the *RNAscope™* Fluorescent Multiplex kit (Cat# 320850, ACDBio, Newark, CA). The following probes were used: Hs-CD9-C3 (Cat# 430671-C3), Hs-PPP1R27-C2 (Cat# 855251-C2), Hs-LRRC10B-C3 (Cat# 855261-C3), Hs-SYT14 (Cat# 855271), Hs-ANXA2-C2 (Cat# 855281-C2), Hs-VSIG10L2-C2 (Cat# 855291-C2), Hs-ADAM11-C3 (Cat# 855311-C3), Hs-CRABP1-C2 (Cat# 855321-C2), Hs-KCNH6-C2 (Cat #855331-C2). Experiments were carried out according to the manufacturer’s instructions (Tissue prep: 320535-TN 11022018, Detection: ACD 320293-UM 03142017), with the following modifications for some tissues to optimize signal quality: boiling time 2.5-5 minutes, protease time 30 min-1 hour, protease temperature 22-40°C.

### Scanning electron microscopy (SEM)

Utricle samples were isolated in washing buffer (0.05 mM HEPES buffer pH 7.2, 10 mM CaCl2, 5 mM MgCl2, and 0.9% NaCl), the otolithic membrane was removed with a brush, and fixed in 4% PFA in 0.05 mM HEPES Buffer pH 7.2, 10 mM CaCl2, 5 mM MgCl2, and 0.9% NaCl for 30 min at RT. The samples were fixed in 2.5% glutaraldehyde and 4% PFA in 0.05 mM HEPES Buffer pH 7.2, 10 mM CaCl2, 5 mM MgCl2, and 0.9% NaCl overnight at 4°C, then washed, dehydrated in ethanol (30%, 75%, 100%, and 100%, 5 min incubation) and processed to the critical drying point using Autosamdri-815A (Tousimis). Samples were mounted on studs using silver paint and coated with 5 nm of Palladium (sputter coater EMS150TS; Electron Microscopy Sciences). Samples were imaged at 5 kV on a FEI Magellan 400 XHR Field Emission Scanning Electron Microscope at the Stanford Nano Shared Facilities.

### Cell quantification and statistical analysis

Using utricles from vestibular schwannoma patients and organ donors, cells were quantified from z-stack images of 25,600 µm^2^, then normalized to 10,000 µm^2^ using Image J64 unless otherwise stated. Images were taken from 1-8 representative areas from the whole sensory epithelium for analyses. For cadaveric utricles, cells were quantified from images of 10,000 µm^2^. Statistical analyses were conducted using Microsoft Excel (Microsoft) and GraphPad Prism 7.0 software (GraphPad). Ordinary one-way ANOVA, Tukey two-way ANOVA and two-tailed, unpaired student’s t-tests were used to determine statistical significance. p<0.05 was considered significant. Data shown as mean±S.D.

### Single cell isolation

Utricles from vestibular schwannoma patients or organ donors were placed in DMEM/F12 (Thermo Fisher Scientific/Gibco, 11-039-021) with 5% FBS during transport to the laboratory. Tissues were then washed twice in DMEM/F12 and any debris and bone microdissected away. The whole utricle was then incubated with thermolysin (0.5 mg/mL; Sigma Aldrich, T7902) for 45 min at 37°C prior to mechanical separation of the sensory epithelium from underlying stroma. Next, the sensory epithelium was digested using Accutase (Thermo Fisher Scientific, 00-4555-56) for 20 min at 37°C and single cell suspension was obtained by trituration using a 1 ml pipette. Single cell suspension was achieved using a 40 μm filter. DMEM/F12 media with 5% FBS was used for this step. Cells were then centrifuged at 300 rcf for 5 min at 4°C. The supernatant was removed and cells were resuspended in media. Number of cells per microliter was quantified using a hemocytometer.

The single cell suspension was then loaded onto a 10x Chromium controller (10x Genomics, Pleasanton, CA) using manufacturer’s recommended protocols at the Stanford Functional Genomics Facility. Following cDNA amplification and library prep, the sample was loaded onto an Illumina HiSeq4000 for sequencing. Two vestibular schwannoma samples were separately processed as described above using two separate 10x Genomics captures and subsequently sequenced in separate lane. One human organ donor sample was also processed as above and the sample was split into two 10x Genomics capture runs and sequenced on two separate lanes.

### Data Preprocessing and Quality Control

Standard 10x Genomics *CellRanger* pipeline was utilized for demultiplexing, alignment, and quantification. Filtered count matrices were then loaded into R to create a *SeuratObject*. We used the *DoubletFinder* method to identify and exclude doublets from four separate 10x captures with a total of 1094 doublets detected using default parameters (McGinnis et al., 2019). Cells with at least 300 features and with less than 20% mitochondrial genes were kept with 735 cells in total excluded based on these criteria. The vestibular schwannoma *SeuratObjects* were then combined, and the organ donor *SeuratObjects* were combined.

### Normalization and Data Integration

Using *Seurat*’s default parameters, the joined data was normalized using *SCTransform* method with the 3,000 most variable genes selected for further downstream analysis. Data were integrated using the ‘IntegrateDat’ function in *Seurat* v3 (Stuart et al., 2019). The integrated data object was then scaled using ‘ScaleDat’ function in *Seurat*.

### Cell Clustering

Following principal component analysis (‘RunPC’) with consideration of 100 principal components, the JackStraw method was used to identify the top 20 most significant principal components. Initial standard UMAP was used for data visualization, followed by clustering with a resolution of 1.0. The *chooseR* algorithm was used to identify the optimal resolution of 1.0 for cell clustering (Patterson-Cross et al., 2021). Given that UMAP representation loses global structure, we chose to represent our data using the previously described Similarity Weighted Non-negative Embedding (SWNE) method for preserving both local and global data structure (Parekh et al., 2018; Wu et al., 2018). The SWNE *Seurat* wrapper function was used per default parameters.

Cell annotation was carried out based on marker genes as described in the literature (Bry et al., 2014; Hammond et al., 2019; Sole-Boldo et al., 2020; Summers et al., 2020; Wilms et al., 2016). In order to focus our analysis on the sensory epithelium, we excluded certain cell clusters (dark cells, stromal cells, melanocytes, vascular cells, roof cells, macrophages, and immune cells). The remaining cells were then reclustered and used for further downstream analysis (Figure S2).

### Differential Gene Expression

The Wilcoxon rank sum test from *Seurat* (Butler et al., 2018; Stuart et al., 2019) was used for identification of differentially expressed genes among the different cell groups with default parameters that utilized the Bonferroni correction for false-discovery (FDR < 0.01). Cell clusters were defined based on known marker genes for each cluster. Only positive marker genes were kept in the differential gene expression analysis. When comparing two groups of cells (Figures 2, 3, S3), we used pseudobulk analysis using the *DESeq2* algorithm. Each 10x capture was considered as a separate run, generating 4 replicates that were used for pseudobulk analysis purposes. For example, there are hair cells from two organ donors and two vestibular schwannoma derived cells that were compared generating a robust differential gene expression comparison. For generation of volcano plots, *DESeq2* results were plotted using *EnchancedVolcano* with indicated thresholds.

### Trajectory Analysis

To identify candidate hair cell precursors, we focused our analysis on the sensory epithelial cells and known hair cell and supporting cell genes. *Seurat* clustering identified seven supporting cell subtypes, two types of hair cells, and one group of putative hair cell precursors. We used *Slingshot* to connect the cell states on the reduced dimensions as visualized by the SWNE plot (Figure S5A) (Street et al., 2018). The reduced dimensions as captured by SWNE was used to build this trajectory using the *Slingshot* algorithm (Street et al., 2018) (Figure 5A, S5A). We defined the starting point as cluster 7, and end points as the two hair cell types. Unbiased lineage trajectory modeling identified four different lineages (Figure S5B) using the default 6 knots within the *tradeSeq* package (Van den Berge et al., 2020). We chose to focus our analysis on the two lineages that generated type I and type II hair cells (Figure 5A). A generalized additive model (GAM) was then used to fit the dynamic expression of each gene along each lineage using the ‘fitGAM’ function in *tradeSeq*. Dynamically expressed genes were identified along each lineage using the ‘AssociationTest’ function. Significantly associated genes were determined using an FDR ≤ 0.01. The ‘diffEntTest’ function in *tradeSeq* identified genes with dynamic expressions that significantly diverge following a common test expression level (Figure 5D). The ‘plotSmoothers’ function allowed the plotting of expression of select individual genes along pseudotime.

### RNA Velocity Analysis

We used the *CellRanger* output *bam* files to identify spliced and unspliced mRNA reads using the *velocyto* command-line (La Manno et al., 2018). The generated *loom* files were imported into a *Python* Jupyter notebook for quantification of spliced to unsliced reads (82% spliced and 18% unspliced in our dataset). They dynamic model algorithm was followed to generate the RNA velocities for each cell (Bergen et al., 2020) projected onto the SWNE reduced dimensions from the analysis above.

## Pathway analysis

Genes that passed *tradeSeq* association test, which assesses significant changes in gene expression as a function of pseudotime, for either type I or type II hair cell lineages with an FDR≤0.01, were uploaded into Ingenuity Pathway Analysis (IPA, QIAGEN Inc., https://digitalinsights.qiagen.com/IPA) for core analysis (Kramer et al., 2014). IPA was performed to identify canonical pathways that are most significant to type I and type II hair cell lineages. Gene Ontology was performed using DAVID to reveal significant biological processes (Huang da et al., 2009a, b).

## Author contributions

T.W., S.E.B., D.K.H., G.S.K., I.A.L., N.G., T.A.J., A.G.C. designed experiments, D.K.H., G.S.K., Z.N.S., K.A.A., Y.V., P.S.M., N.H.B., R.K.J., A.I., I.A.L., T.A.J. procured tissues, T.W., A.H.L., N.P., D.W., S.E.B., N.G., T.A.J., I.A.L., M.S., A.G.C. performed experiments, T.W., A.H.L., S.E.B., G.S.K., P.A.J., N.G., S.H., T.A.J., A.G.C. analyzed data, T.W., A.H.L., S.E.B., P.J.A., N.G., T.A.J., A.G.C. prepared the manuscript.

## Acknowledgements

We thank for our laboratory for insightful comments on the manuscript, J. Alyono, J. Oghalai, W. Dong, J. Burns, D. Ellwanger for excellent technical support, Y. Ma for statistical analyses, H. Bellen for sharing reagents, and A. Salehi and staff at Donor Network West and Stanford Otolaryngology surgeons for assistance with tissue procurement. This work was supported by NSF ECCS-2026822 (the Stanford Nano Shared Facilities), S10OD018220, S10OD021763 (Stanford Functional Genomics Facility), the Lucile Packard Foundation for Children’s Health, Stanford NIH-NCATS-CTSA UL1 TR001085, Child Health Research Institute of Stanford University, NIDCD/NIH R21DC015879, National Natural Science Foundation of China 82071056 (T.W.), Stanford University Medical Scholars Research Program, Howard Hughes Medical Institute Medical Fellows Program, NIDCD/NIH F30DC015698 (Z.N.S.), U24DC015910 (A.I. and I.A.L.), R01DC016409, R21DC019457 (N.G.), K08DC019683, Hearing Research, Inc. (T.A.J.) and R01DC013910, T32DC015209, U24DC020857, Department of Defense MR130316, Akiko Yamazaki and Jerry Yang Faculty Scholar Fund, and California Initiative in Regenerative Medicine RN3-06529 (A.G.C.), and generous support by the Yu, Ogawa, and Donoho families and the Bill and Susan Oberndorf Foundation.

## Supplementary items

Figure S1 (to Figure 1) Surgical approaches to procure and quality control for human utricles

Figure S2 (to Figure 2) Strategies to subset and validate enriched genes in human hair cells and supporting cells

Figure S3 (to Figure 3) Genes enriched in hair cell and supporting cell subtypes

Figure S4 (to Figure 4) Degenerating hair cells in utricles from vestibular schwannoma (VS) patients.

Figure S5 (to Figure 5) Trajectory analysis

Figure S6 (to Figure 6) Characteristics of hair cell precursors in organ donor and vestibular schwannoma utricles

Figure S7 (to Figure 1) Expression patterns of genes associated with hearing loss and vestibular dysfunction

Table S1 (to all figures) Patient demographic and medical information

Table S2 (to Figure 1) Differential gene expression of all cells

Table S3 (to Figures 2 and S2) Differential gene expression of sensory epithelial cells

Table S4 (to Figure 2) Hair cell versus supporting cell markers

Table S5 (to Figure 3) Hair cell subtype genes

Table S6 (to Figure S3) Supporting cell subtype genes

Table S7 (to Figures 4 and S4) Quantification of hair cells and supporting cells

Table S8 (to Figure 5) Type I hair cell lineage associated genes

Table S9 (to Figure 5) Type II hair cell lineage associated genes

Table S10 (to Figure 5) Type I hair cell lineage associated pathway analysis

Table S11 (to Figure 5) Type II hair cell lineage associated pathway analysis

Table S12 (to Figure 5) Type I hair cell lineage GO terms

Table S13 (to Figure 5) Type II hair cell lineage GO terms

Table S14 (to Figure 6) Type I hair cell lineage transcription factors

Table S15 (to Figure 6) Type II hair cell lineage transcription factors

Table S16 (to all Figures) Key resource table

